# Identification of insertion sites for the integrative and conjugative element Tn9*16* in the *Bacillus subtilis* chromosome

**DOI:** 10.1101/2025.01.28.635231

**Authors:** Emily L. Bean, Janet L. Smith, Alan D. Grossman

**Affiliations:** Department of Biology, Massachusetts Institute of Technology, Cambridge, MA 02139

**Keywords:** Horizontal gene transfer, mobile genetic elements, integrative and conjugative element, conjugative transposon, Tn*916*, *Bacillus subtilis*, integration sites, nucleoid associated protein, *rok*

## Abstract

Integrative and conjugative elements (ICEs) are found in many bacterial species and are mediators of horizontal gene transfer. Tn*916* is an ICE found in several Gram-positive genera, including *Enterococcus*, *Staphylococcus*, *Streptococcus*, and *Clostridum*. In contrast to the many ICEs that preferentially integrate into a single site, Tn*916* can integrate into many sites in the host chromosome. The consensus integration motif for Tn*916*, based on analyses of approximately 200 independent insertions, is an approximately 16 bp AT-rich sequence. Here, we describe the identification and mapping of approximately 10^5^ independent Tn*916* insertions in the *Bacillus subtilis* chromosome. The insertions were distributed between 1,554 chromosomal sites, and approximately 99% of the insertions were in 303 sites and 65% were in only ten sites. One region, between *ykuC* and *ykyB* (*kre*), was a ‘hotspot’ for integration with ∼22% of the insertions in that single location. In almost all of the top 99% of sites, Tn*916* was found with similar frequencies in both orientations relative to the chromosome and relative to the direction of transcription, with a few notable exceptions. Using the sequences of all insertion regions, we determined a consensus motif which is similar to that previously identified for *Clostridium difficile*. The insertion sites are largely AT-rich, and some sites overlap with regions bound by the nucleoid-associated protein Rok, a functional analog of H-NS of Gram-negative bacteria. Rok functions as a negative regulator of at least some horizontally acquired genes. We found that the presence or absence of Rok had little or no effect on insertion site specificity of Tn*916*.

## Introduction

Integrative and conjugative elements (ICEs), also called conjugative transposons, are a type of self-transmissible mobile genetic element that facilitates horizontal gene transfer and contributes to bacterial evolution (Wozniak and Waldor, 2010; Bellanger *et al*., 2014; Johnson and Grossman, 2015; Delavat *et al*., 2017). ICEs typically carry accessory (cargo) genes that benefit the host cell, including genes that confer antibiotic resistances, pathogenic or symbiotic determinants, or alternative metabolic pathways (Frost *et al*., 2005; Treangen and Rocha, 2011; Johnson and Grossman, 2015). Typically, when an ICE is integrated in a host genome, the ICE genes needed for conjugation are not expressed. Either stochastically, or upon some environmental stimulus, an ICE can excise from the chromosome and the genes needed for conjugation are expressed. The element DNA can then be nicked and unwound to generate ssDNA that can then be transferred through the element-encoded conjugation machinery, a Type IV secretion system (T4SS) into a recipient cell. Once in the recipient, the ICE DNA eventually integrates into the chromosome of the new host to generate a stable transconjugant (for reviews, see (Wozniak and Waldor, 2010; Johnson and Grossman, 2015)).

Tn*916* was the first described ICE and was discovered because of its ability to transfer tetracycline resistance between isolates of *Enterococcus faecalis* (Franke and Clewell, 1981a; Franke and Clewell, 1981b). Since its initial discovery, Tn*916* and Tn*916-*like elements have been found in several Gram-positive host species, including *E. faecalis*, *Clostridium difficile, Staphylococcus aureus,* and *Streptococcus pneumonia* (Fitzgerald and Clewell, 1985; Clewell *et al*., 1985; Clewell and Flannagan, 1993; Roberts and Mullany, 2009; Roberts and Mullany, 2011; Santoro *et al*., 2014; Sansevere and Robinson, 2017). Additionally, Tn*916* is functional and its biology has frequently been studied in *B. subtilis* (e.g. (Ivins *et al*., 1988; Scott *et al*., 1988; Roberts *et al*., 2003; Wright and Grossman, 2016)).

ICEs encode integrases (recombinases) that catalyze integration into and excision out of a host chromosome. Many ICEs have a preferred integration (attachment) site in the chromosome, often in a conserved and essential gene (Burrus and Waldor, 2004). If the preferred site is absent, secondary attachment sites can be used (Wang *et al*., 2000a; Wang *et al*., 2006; Lee *et al*., 2007; Menard and Grossman, 2013), analogous to temperate phages that can utilize secondary attachment sites (e.g. Shimada et al., 1972) and non-conjugative transposons, e.g., Tn7 (reviewed in (Craig, 1991)).

Some ICEs, including those in the Tn*916* family, are promiscuous in target site selection and can integrate into many sites in a host chromosome (Clewell *et al*., 1988; Scott *et al*., 1994b; Cookson *et al*., 2011a; Mullany *et al*., 2012). Tn*916* integrates into AT-rich regions in a host chromosome (Clewell *et al*., 1988; Scott *et al*., 1994a; Cookson *et al*., 2011b; Mullany *et al*., 2012). To date, the most extensive characterization of Tn*916* integration site selection was completed in the context of *C. difficile* clinical isolates. Based on the sequences of approximately 200 independent Tn*916* insertions the consensus insertion motif was defined as the AT-rich sequence 5’-TTTTTA[AT][AT][AT][AT]AAAAA (Mullany *et al*., 2012). A similar site is used in *Butyrivibrio proteoclasticus* (Cookson et al., 2011).

Here, we describe the sequence of sites of Tn*916* integration in the *B. subtilis* chromosome from ∼10^5^ independent Tn*916* transconjugants. The ∼10^5^ insertions were distributed between 1,554 chromosomal sites, with widely varying frequencies in each site. Ninety-nine percent of insertions were in 303 sites, 65% were in only ten sites, and 22% were in a single (hotspot) site. In almost all of the top 99% of sites, Tn*916* was found with similar frequencies in both orientations relative to the direction of DNA replication and transcription (for insertions in open reading frames), with a few notable exceptions. Insertions were centered around an AT-rich consensus motif that is similar to that previously described for Tn*916* insertions in *C. difficile* (Mullany *et al*., 2012) and *B. proteoclasticus* (Cookson et al., 2011).

We also determined the insertion site specificity of Tn*916* in cells missing the nucleoid associated protein Rok. Rok binds to AT-rich regions of the chromosome (Smits and Grossman, 2010; Seid *et al*., 2017) and is analogous to H-NS (reviewed in (Grainger, 2016), found in many Gram-negative bacteria. We did not detect a global effect of *rok* on insertion site specificity of Tn*916*, especially at the more abundant insertion sites.

## Results and Discussion

### Rationale and overview

We aimed to identify Tn*916* integration sites in the *B. subtilis* chromosome and to determine if the nucleoid binding protein Rok substantially altered the integration site preferences. Based on previous analyses of Tn*916* insertions in other organisms (e.g. Cookson et al., 2011; Mullany et al., 2012; Scott et al., 1994), we anticipated that insertions in *B. subtilis* would also be in AT-rich regions with a consensus similar (or identical) to that found in other species. These previous findings motivated our experiments to test the effects of Rok, a non-essential nucleoid-associated protein that preferentially binds AT-rich regions of the chromosome and can repress expression of some horizontally acquired genes (Smits and Grossman, 2010; Seid *et al*., 2017), analogous to the roles of H-NS in Gram-negative bacteria (Dorman, 2007; Navarre *et al*., 2007; Fang and Rimsky, 2008; Smits and Grossman, 2010; Seid *et al*., 2017). We used high-throughput sequencing to identify Tn*916* insertion sites in the *B. subtilis* genome and determined if Rok had any major effects on utilization of these sites.

### Libraries of Tn*916* insertions

We made libraries of approximately 10^5^ independent Tn*916* insertions in both wild type and Δ*rok B. subtilis* cells. Briefly, a Tn*916* donor (host) strain (ELC43) containing a copy of Tn*916* integrated near the *yufK* start codon was crossed with two recipient strains that did not already contain a copy of Tn*916* (*rok+* strain ELC38 and Δ*rok* strain ELC42) and transconjugants that obtained Tn*916* were selected (Methods). The conjugation efficiencies using wild type and *Δrok* recipients were both ∼0.02% transconjugants per donor. Transconjugants from multiple matings were recovered from petri plates, pooled, and then grown in LB medium to stationary phase before harvesting genomic DNA to map Tn*916* insertion sites by high throughput sequencing.

### Identification of insertion sites

We identified the Tn*916* insertion (integration) sites in transconjugants using a transposon-directed insertion sequencing method (TraDIS) (Langridge *et al*., 2009; Potter and Luo, 2010; Mullany *et al*., 2012; Barquist *et al*., 2016). Briefly, genomic DNA from the pooled transconjugants (*rok+* and *Δrok*, separately) was randomly sheared, adaptors were annealed to the ends of the fragments, and the junctions between the left end of Tn*916* (as drawn in Fig 1A) and the chromosome were amplified by PCR. A Tn*916*-specific primer that hybridizes between *orf24* and *attL* was used for sequencing (Fig 1B, Methods). The relative number of sequence reads for a given site reflects the relative number of insertions in that site after growth of the cells, assuming that all other reactions are of equal efficiencies for each insertion. Approximately 87% of the sequence reads from each pool (*rok*+ and Δ*rok*) mapped to the *B. subtilis* chromosome and five percent of the total reads were from the excised circular form of Tn*916*.

**Fig 1.**
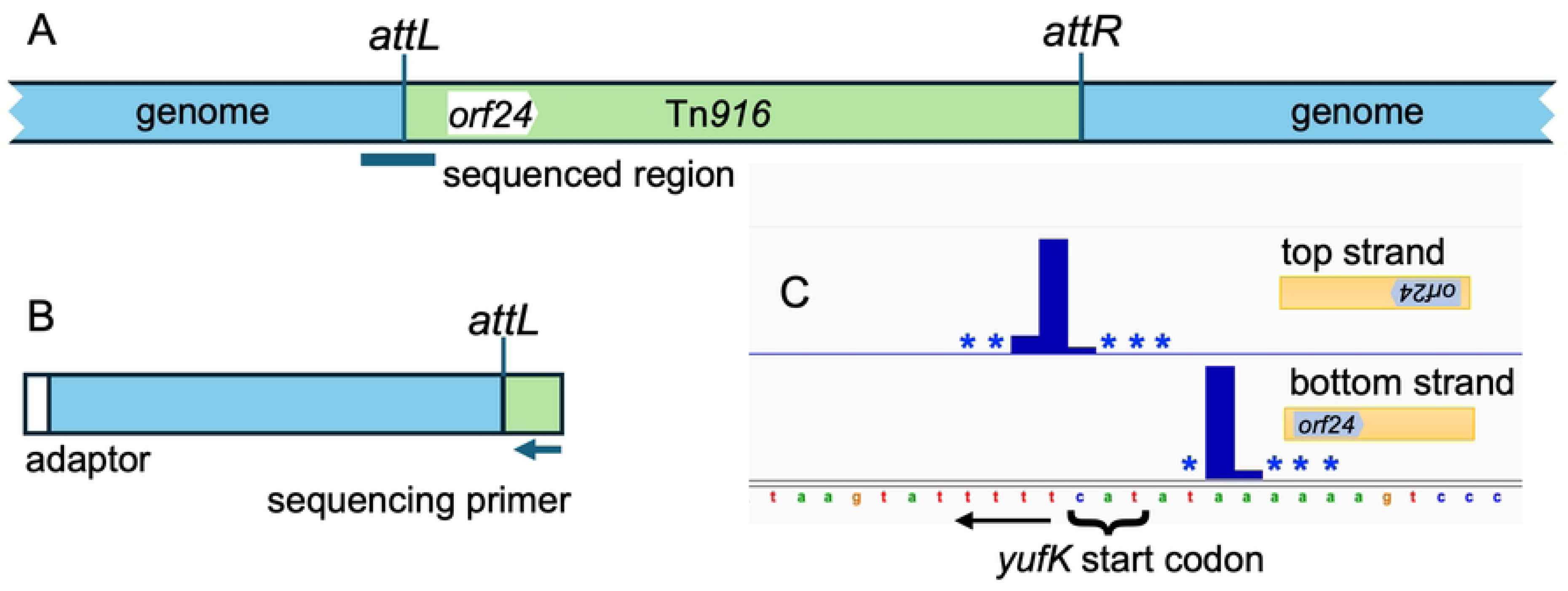
Schematic of Tn*916* and identification of insertion sites. **A.** A generic insertion of Tn*916* (green) in the chromosome (blue) is shown, including *attL*, *attR*, and *orf24* (the first open reading frame within Tn*916*). DNA for sequencing was amplified from the left end of Tn*916* (as drawn) extending into the chromosome, as indicated below the insertion. **B.** Fragments that span the insertion site at the left end of Tn*916* were amplified from the library of transconjugants, resulting in fragments approximately 300 bp in length. Amplification was carried out using an adaptor (shown in white) for the chromosomal end together with a primer that anneals within Tn*916* (see Methods for details). The sequencing primer binds between *attL* and *orf24*. Reads that map to the bottom DNA strand correspond to insertions in the orientation drawn in panel A; reads mapping to the top strand are in the opposite orientation, *i.e.*, with *orf24* the right. **C.** Tn*916* insertions at *yufK*. Insertions occur in both orientations. Insertions with *orf24* to the left had DNA sequence reads on the bottom strand and insertions with *orf24* to the right had sequence reads on the top strand. Insertions were at multiple adjacent nucleotides and relative frequencies are indicated by the bar plots, with an asterisk indicating an insertion position used at a frequency too low to plot to scale.

Insertions at any given genomic locus were often observed in both orientations and at multiple adjacent or nearby bases (e.g., Fig 1C). Because these nearby insertions are associated with the same insertion motif, we grouped them together into insertion sites (see Methods). For wild-type cells, insertions occurred at a total of 5377 nucleotide positions and clustered into 1554 insertion sites (Methods; S1 Table). The frequency of insertions in different sites varied over a wide range in both *rok*+ and Δ*rok* strains (Fig 2 and S1 Table).

**Fig 2.**
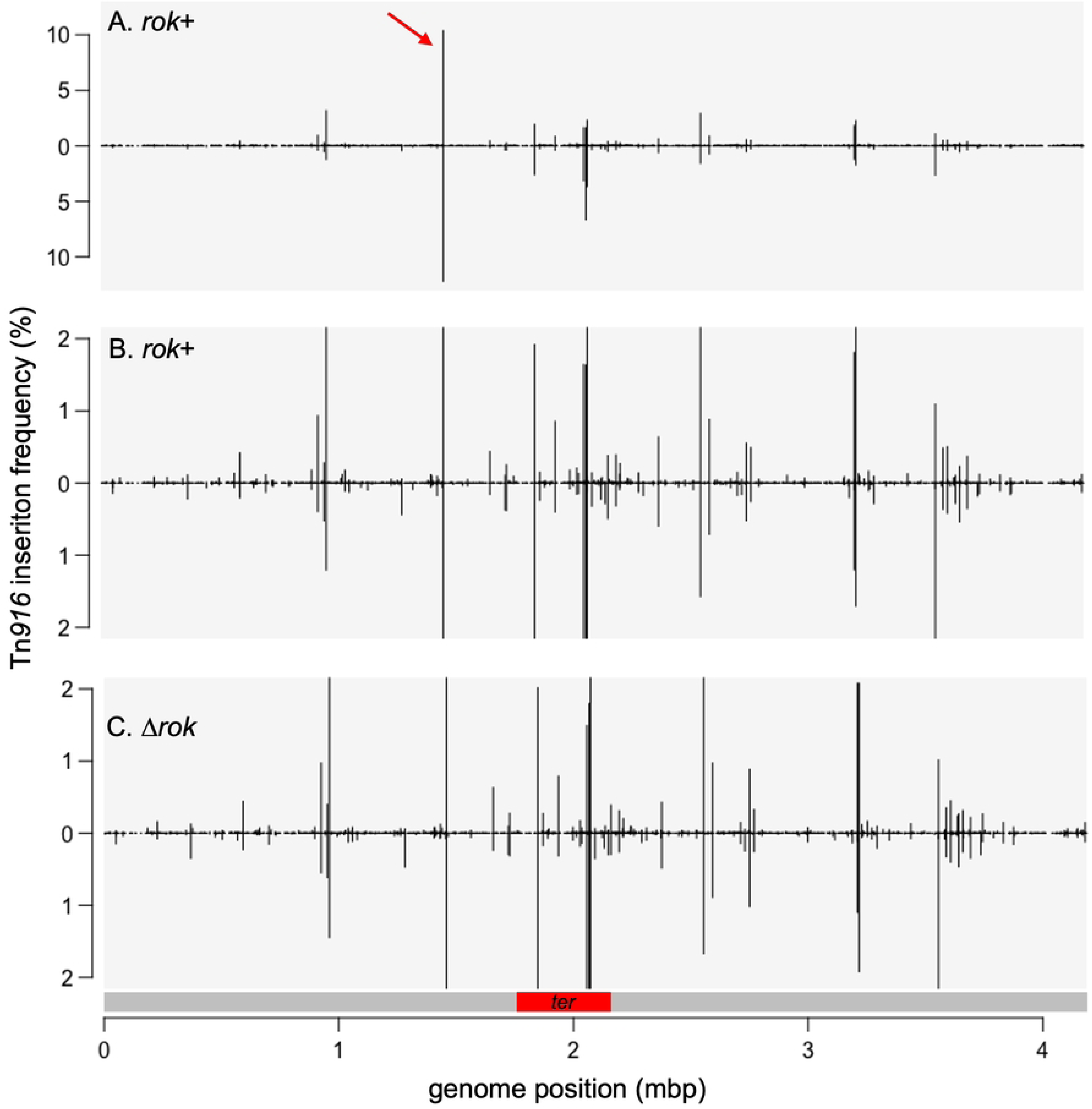
Relative frequencies of Tn*916* insertions throughout the chromosome. The relative frequency of Tn*916* insertions along the chromosome with the origin of replication at the left end and the terminus region near the middle are presented. Insertions on the top strand are shown above the line and those on the bottom strand below the line. The linear depictions of the chromosome begin and end at the origin of replication and the red bar below (C) indicates the *ter* region. **A, B**) Insertions from mating Tn*916* into wild type strain (ELC38). Data are the same in each panel, but the scale in (B) is different to accentuate insertions in sites used at a low frequency. The red arrow in (A) indicates the insertion hotspot between *ykuC* and *ykyB* that contains ∼22% of the total insertions. **C**) Insertions from mating Tn*916* into the Δ*rok* strain (ELC42). Data are presented on the same scale as in B for comparison.

We found that 99% of the Tn*916* insertions in wild type cells were in 303 sites (S1 Table). Of these 303 sites, 59% were outside of annotated open reading frames (intergenic), although only ∼12% of the genome is intergenic. Taking into account the frequency of insertions in each site, 75% of the insertions in these 303 sites were in intergenic regions. This bias is likely driven, in part, by the AT-rich nature of these regions (see below). A similar bias was observed for Tn*916* insertions in *C. difficile* (Mullany *et al*., 2012).

### The Tn*916* insertion site motif

We defined a consensus sequence motif for insertion of Tn*916* in the *B. subtilis* chromosome using MEME (Bailey and Elkan, 1994a). We submitted 61-bp of DNA sequence surrounding the 303 most frequently used sites in wild type cells. The resulting consensus motif contained a palindromic region of T- and A-rich stretches separated by six base pairs that are AT-rich but less well conserved (Fig 3A). This motif has more sequence information, but is virtually the same as the consensus motif identified in other organisms (e.g. Cookson et al., 2011; Mullany et al., 2012; Scott et al., 1994).

**Fig 3.**
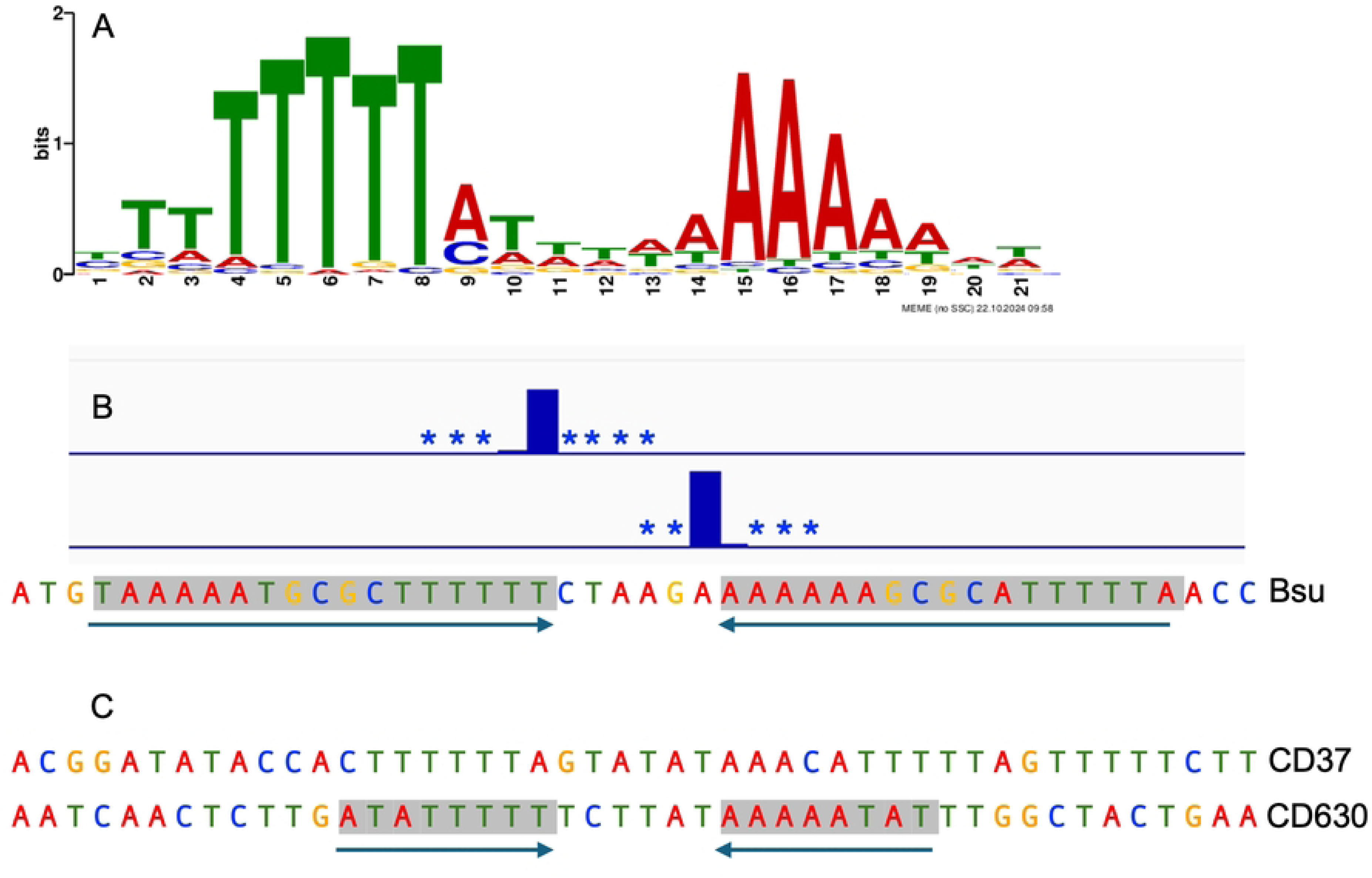
The consensus Tn*916* insertion site motif. **A**) The consensus sequence logo for the top 303 Tn*916* insertion sites in wild type cells. The consensus sequence is similar to that determined for Tn*916* insertions in other organisms (Cookson et al., 2011; Mullany et al., 2012) **B)** Tn*916* insertions in the hotspot between *ykuC* and *ykyB* (*kre*), where 22% of insertions occur. Insertions in the reverse (top strand) or forward (bottom strand) are indicated by bar plots, with additional insertion positions present at frequencies too low to plot to scale indicated by asterisks. A 17-bp perfect inverted repeat is indicated by the black arrows and gray highlighting. **C)** Sequence of the Tn*916* insertion hotspot in *C. difficile* strain CD37 (Mullany *et al*., 1991; Wang *et al*., 2000) and in *C. difficile* strain 630 (Mullany *et al*., 2012). Shading and arrows indicate the 8 bp inverted repeat in the hotspot in CD630.

### Hotspots for Tn*916* insertions

Remarkably, 65% of the independent Tn*916* insertions were in ten sites (Table 1; Fig 2A), and 22% were in a site between *ykuC* and *ykyB* (*kre*). This bias for insertions in the *ykuC*/*ykyB* site was not because the Tn*916* from donor strain had been integrated in this site. The strain used as a donor (ELC43) had a single copy of Tn*916* at the very 5’-end of *yufK* (Wright and Grossman, 2016)). Tn*916* integrated at this site in ∼3% of transconjugants. We suspect that the bias for insertions in the *ykuC*/*ykyB* site could be due to the presence of a 17 bp inverted repeat that flanks the nucleotides where insertions occur (Fig 3B), and none of the other insertion sites had a 17 bp inverted repeat. We have not tested experimentally the importance of this inverted repeat on insertion frequency.

**Table 1.**
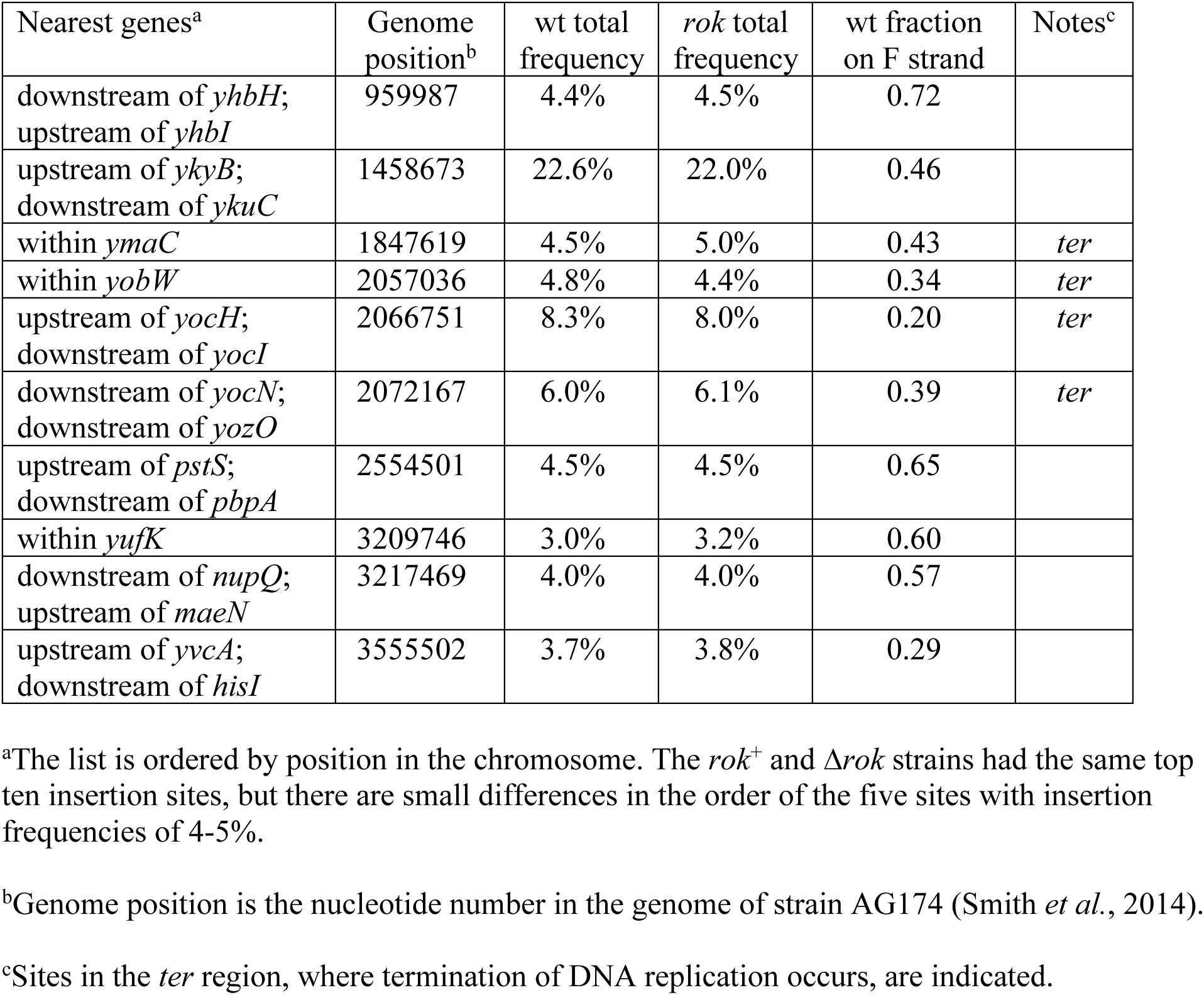
Top ten Tn*916* insertion sites in *B. subtilis*.

The high frequency (∼22%) of Tn*916* insertions in a single site in the *B. subtilis* chromosome is intriguing when considered with the previously reported finding that Tn*916* has one highly preferred integration site in the *C. difficile* strain CD37 (eight independent integrations in this site, one transconjugant from each of eight different conjugation experiments), despite the presence of multiple potential integration sites in the chromosome (Mullany *et al*., 1991; Wang *et al*., 2000). This hotspot is located downstream of *buk2* and does not have an inverted repeat. In *C. difficile* strain 630 there are a range of insertion frequencies at various sites, with insertions occurring most frequently near the 3’ end of the *mgtA* coding sequence (Mullany *et al*., 2012). This insertion site is immediately flanked by a perfect 8 bp inverted repeat. Although this is much shorter than the 17 bp inverted repeat at the *B. subtilis* hotspot, it is consistent with the possibility that inverted repeats can increase insertion frequency.

Overall, our results are consistent with those previously reported (Mullany *et al*., 1991; Wang *et al*., 2000b; Mullany *et al*., 2012), and together indicate that insertion site specificity is determined by a combination of sequence and other factors that influence DNA conformation (Mullany *et al*., 1991; Wang *et al*., 2000b; Mullany *et al*., 2012). Because the Tn*916* recombinase (Int) has a preference for DNA sites with a static bend (Lu and Churchward, 1995), the preferred sites in different organisms likely reflect that local DNA structure in combination with the sequence. Static bends are associated with polyA tracts (Haran and Mohanty, 2009), which are a feature in the Tn*916* insertion motif, but additional sequence elements and DNA binding proteins may enhance the formation of static bends in these regions.

In addition to reflecting preferences for specific DNA sequences and structures, the frequency at which insertions in each site are recovered is also influenced by the selective pressures on the insertion mutants during growth. The vast majority of genes are not expected to have a significant effect on fitness under the rich media growth conditions used here (Kobayashi *et al*., 2003; Johnson and Grossman, 2014; Peters *et al*., 2016; Koo *et al*., 2017), and the outgrowth used to prepare our library was likely not enough for small differences in fitness to have a large effect of the final population. Thus we believe that the bulk of the differences we observe in the frequencies of insertion in different regions are due to DNA site preferences and not effects of the insertions on fitness.

### Increased Tn*916* insertion frequency in the terminus region of the chromosome

Four of the ten insertion sites that are utilized most frequently are in the ∼10% of the chromosome that comprises the terminus (*terC*) region of replication (Fig 2A and 2B). As a result, this region (indicated by a red bar in Fig 2) has a 3.8-fold higher insertion frequency compared the rest of the chromosome. The matings were done during mid- and late-exponential phase in rich medium, and on average, there are approximately four copies of the origin region for each copy of the terminus region in these growth conditions. This indicates that the inherent preference for the sites in the *terC* region may be stronger than the observed frequencies indicate. These high frequency sites are not coincident with the any of the nine *ter* sites where termination actually occurs. Nor are they coincident with the *dif* site which is in the *ter* region and is used by the site-specific recombinases CodV and RipX to resolve chromosome dimers that can form during replication (Sciochetti *et al*., 2001; Gogou *et al*., 2021). The *C. difficile* strain R20291 may also have an elevated frequency of insertions in the *ter* region (Mullany *et al*., 2012), indicating that there might be a feature of this chromosomal region, separate from the *ter* and *dif* sites, that favors Tn*916* insertions.

### Orientation bias at insertion sites

We analyzed the frequency of insertions on the forward versus reverse strand at each insertion site (Fig 4). For the top 99% of insertion sites (303 sites; black dots in Fig 4), 97% had insertions in both orientations. The insertion sites that had a strong orientation bias (≥4-fold in one orientation) tended to have lower numbers of insertions. We suspect that in many cases, the apparent orientation bias is likely due to the relatively low frequency of insertions in these sites and is not an indication of the properties of the insertion site *per se*. In general the insertion frequency in each orientation at individual sites was similar, especially for more abundant insertion sites. However even among the 10 most abundant sites (Table 1) there were some that had more than a two-fold bias in one orientation, indicating that there might be features of individual sites that drive asymmetric insertion frequencies.

**Fig 4.**
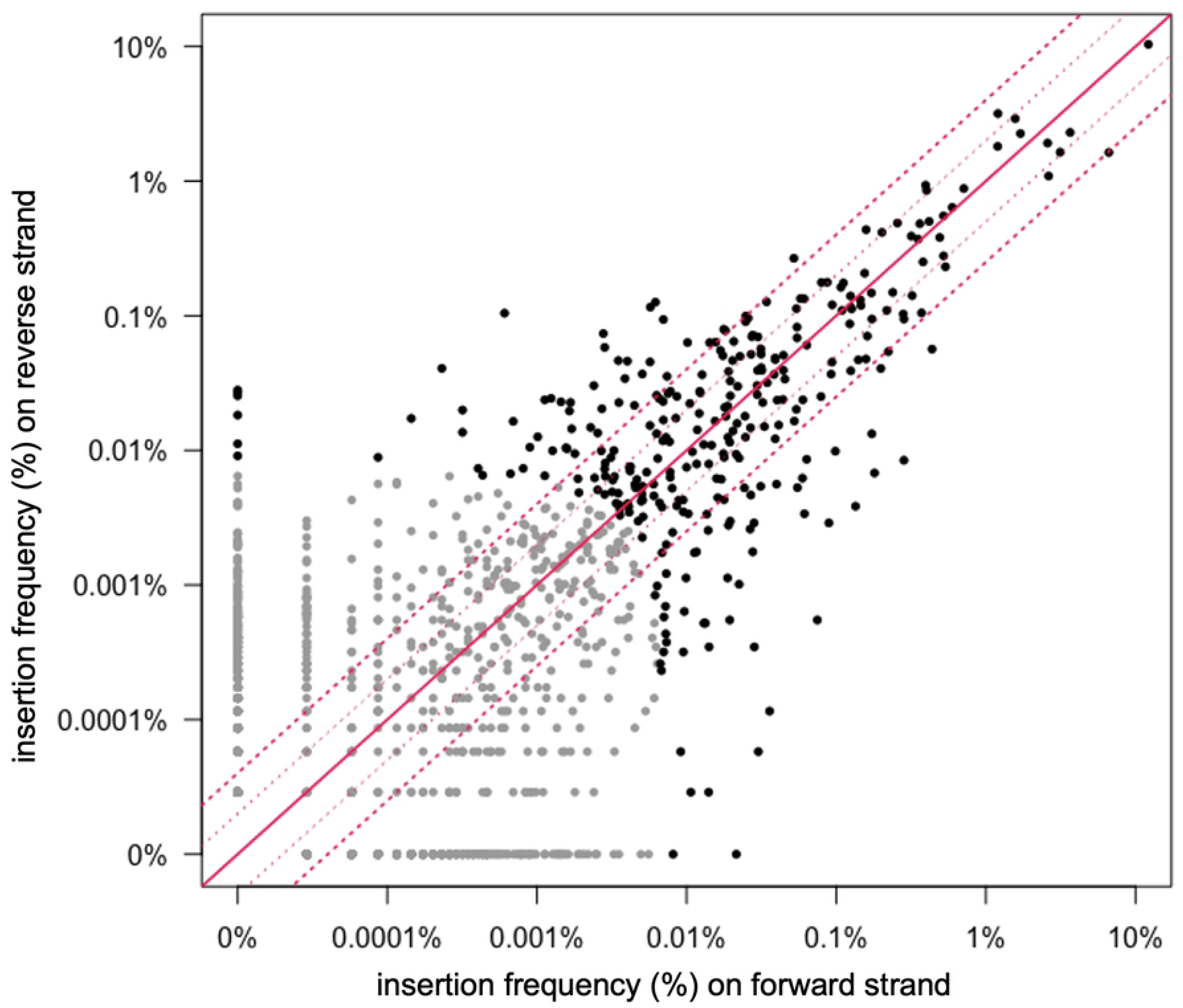
Orientation and frequency of insertions in each site. The frequency of insertions in each orientation in the reverse vs forward directions are plotted for all 1,554 insertion sites in wild type cells. The 303 most frequently used sites are shown in black and the rest are in gray. The solid red diagonal line indicates an equal number of insertions in each orientation. The dotted red lines indicate a 2- and 4-fold difference in abundance.

The orientation of Tn*916* insertions was not strongly influenced by leading- vs lagging-strand DNA replication (Fig 2). In the region from the origin of replication (*oriC*) going clockwise to the beginning of the terminus (*ter*) region, insertions were equally distributed between forward (leading strand) and reverse (lagging strand) orientation, and in the region from *oriC* going counterclockwise to the start of the *ter* region, 52% of insertions had *orf24* oriented codirectionally with leading strand DNA synthesis (Fig 2A and 2B). The *terC* region was not included in this analysis because leading vs. lagging strand synthesis across this region is not clearly demarked.

### Orientation of insertions relative to host transcription

To determine if there was a systematic orientation bias related to host transcription, we analyzed the orientation of Tn*916* insertions in open reading frames. Of the top 303 insertion sites (99% of insertions), 124 were in an open reading frame (S1 Table). We found that 56% of the insertions in these 124 sites were oriented such that *orf24* was codirectional with the host ORF. Based on these results, we conclude that there appears to be no strong systematic bias for insertion orientation in transcribed portions of the chromosome.

Despite the absence of systematic bias, there appeared to be an orientation bias of more than four-fold in nine sites in open reading frames with an insertion frequency of at least 0.1% (Fig 5, Table 2). Three of the sites had a higher frequency of insertions with *orf24* codirectional with transcription of the host *orf*, and six sites had a higher frequency of insertions in the opposite orientation. These biases could be due to random fluctuations or might reflect inherent differences in DNA structure that influences integration into these sites.

**Fig. 5.**
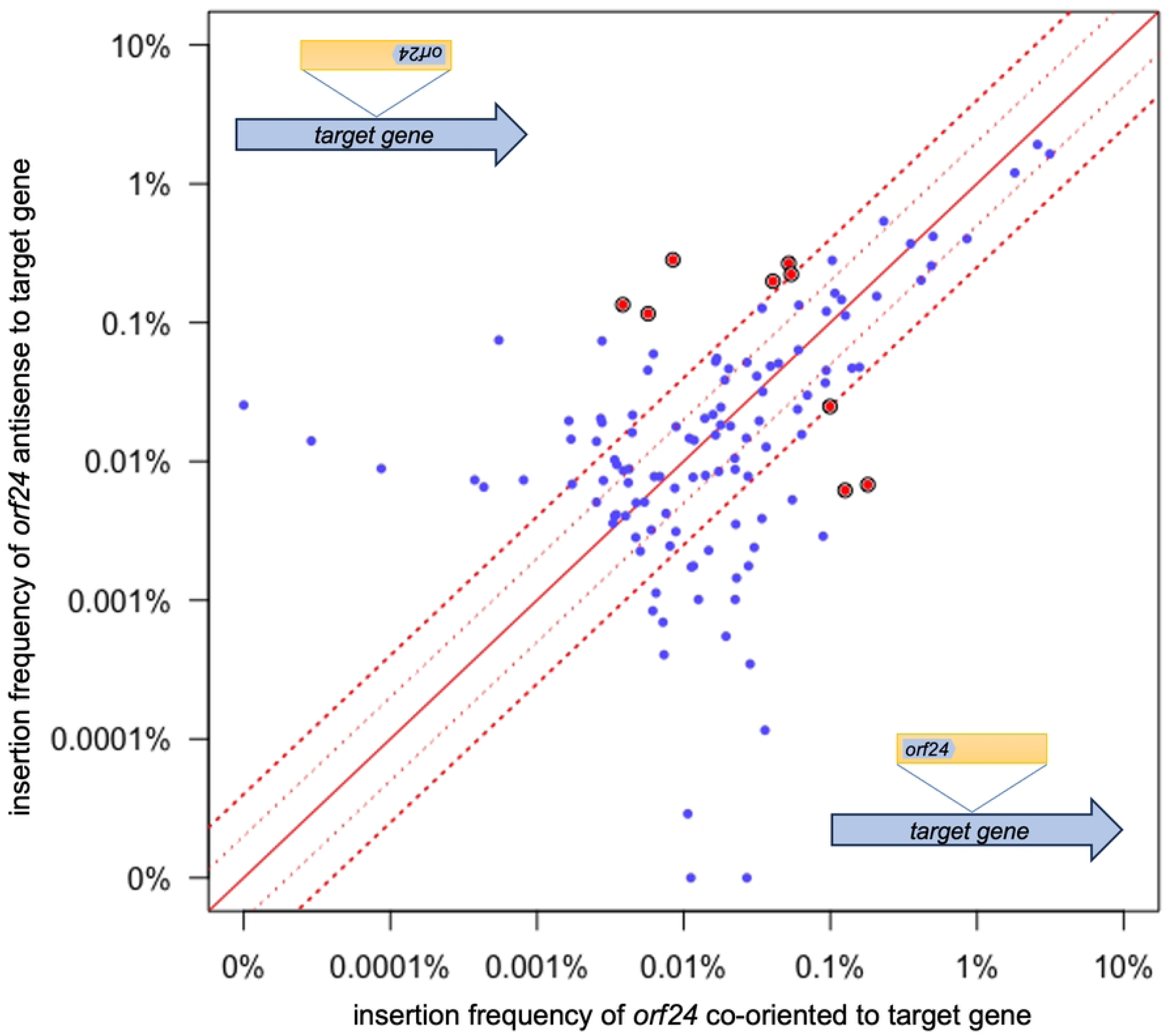
Insertion orientation in sites within ORFs. Of the top 303 insertion sites, 124 are within *orfs*. The frequencies of Tn*916* insertions in each of these sites with *orf24* codirectional (x-axis) or antisense (y-axis) to transcription of the host gene are plotted. The solid red diagonal line indicates an equal number of insertions in each orientation. The dotted red lines indicate a 2- and 4-fold difference in abundance. The filled red circles indicate sites that had ≥4-fold strand bias and an insertion frequency of ≥0.1%. These are also listed in Table 2.

**Table 2.** Insertion sites in open reading frames with at least 4-fold orientation bias.

| Gene | Genome position | Gene strand | wt total frequency<sup>a</sup> | Percent with *orf24* codirectional<sup>b</sup> |
| --- | --- | --- | --- | --- |
| *yheJ* | 1028733 | + | 0.12% | 5% |
| *mtnU* | 1409015 | - | 0.12% | 80% |
| *yosV* | 2130878 | - | 0.28% | 20% |
| *yorM* | 2148167 | - | 0.29% | 3% |
| *yomW* | 2212241 | + | 0.32% | 16% |
| *yugS* | 3188952 | - | 0.24% | 17% |
| *comP* | 3228200 | - | 0.13% | 95% |
| *clsA* | 3735469 | + | 0.19% | 96% |
| *sacT* | 3879050 | - | 0.14% | 3% |

<sup>a</sup>The sites included here have an insertion frequency of at least 0.1% of the total number of insertions and at least 80% of the insertions are in one orientation.
<sup>b</sup>The percent of insertions in the indicated site with *orf24* from Tn916 codirectional with transcription of the host gene.

There might also be selective pressures against insertions in a particular orientation that are driven by transcription, but that are specific to a given locus. For example, if the host gene is regulated by an antiterminator between the promoter and insertion site, there might be read-through of the terminator at the left end of Tn*916*, which would cause a competitive disadvantage due to expression of genes involved in unwinding and rolling circle replication (Wirachman and Grossman, 2024). In addition, transcription from promoters at the right end of Tn*916* (P*orf7*, P*xis*, P*int*) that are directed into the chromosome might cause expression of regions downstream of the insertion site (with *int* distal to the host promoter) or antisense transcription of regions upstream the insertion site (with *int* proximal to the host promoter). If such transcription is deleterious, it would be gene/insertion specific.

### Evaluation of Rok effects on insertion site selection

Rok is a nucleoid-associated protein that binds AT-rich sequences (Smits and Grossman, 2010; Seid *et al*., 2017) and could potentially affect the ability of Tn*916* to integrate into specific sites. We compared the frequencies of Tn*916* insertions in all sites in a *rok* null mutant vs. wild type (*rok*^+^), and also specifically analyzed insertions in sites that are in known Rok binding regions (Seid *et al*., 2017). In general, there were similar insertion frequencies in most sites in *rok*+ and *Δrok* strains (Fig 6), especially in sites with higher insertion frequencies. Thirteen of the insertion sites were in known Rok binding regions (Fig 6, red circles). A few of these had different insertion frequencies in Δ*rok* and *rok^+^* cells, but no consistent pattern was observed as to whether there was a higher insertion frequency in Δ*rok* or *rok^+^* cells. Since the sites that had larger difference in the two strains were sites used at a relatively low frequency, they have a higher chance of differing due to random fluctuation. Based on these analyses, we conclude that Rok does not substantially or systematically affect the insertion site utilization of Tn*916*. We have not determined if Rok affects excision of Tn*916* from the chromosome. If there are any effects on excision, we suspect that they would occur at a small number of sites.

**Fig 6.**
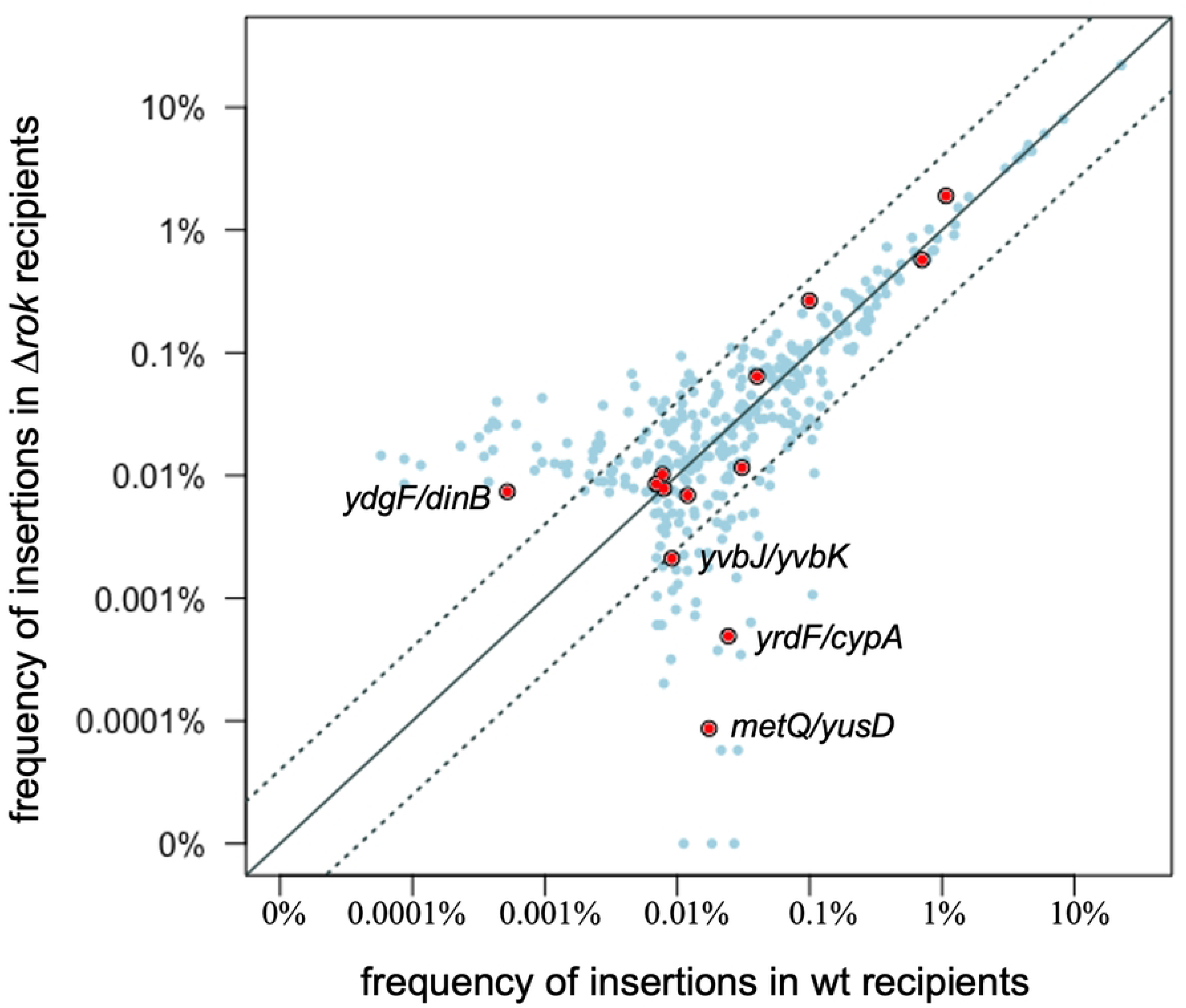
Comparison of insertion frequencies in wild type and Δ*rok* strains. The frequency of insertions in each of the top 99% of sites in the Δ*rok* mutant (302 sites) are plotted versus the frequency in wild type cells (303 site). Filled red circles indicate Rok binding regions previously reported (Seid *et al*., 2016). The solid diagonal line indicates an equal number of insertions in each strain. The dotted lines indicate a 4-fold difference in insertion frequency. The location of the four Rok binding sites that have >4-fold difference in insertion frequency in between the two strains is indicated with the gene names.

### Conclusions and perspectives

Overall, we identified >1900 unique Tn*916* insertion sites in the *B. subtilis* chromosome. We did not observe a systematic influence of Rok on integration site selection. Despite the relatively large number of independent insertions analyzed, the insertion sites were not saturated. It could prove interesting to evaluate the integration site distribution after removing the high-frequency insertion site from the chromosome (between *ykuC* and *ykyB*), and perhaps others, to favor the insertion into the lower-frequency sites. Additionally, the utility of Tn*916* as a potential mutagen has been previously explored and the limitations noted (Ivins *et al*., 1988; Lin and Johnson, 1991; Awad and Rood, 1997; Mullany *et al*., 2012). The preference exhibited in target site selection in *C. difficile* and *B. subtilis* indicates that Tn*916* insertion sites lack the desired randomness and diversity of target sites for random transposon mutagenesis across the chromosome.

One of the most interesting aspects of Tn*916* integration is the presence of integration ‘hotspots’ in different organisms. The sequence of the hotspot in one organism is different from that in another (e.g., *B. subtilis* vs *C. difficile*), even though the same recombinase catalyzes the reaction and the consensus integration site is virtually the same in each organism. Consistent with other analyses, our findings indicate that there are likely multiple factors that contribute to the use of one site preferentially over others, perhaps including genome location and organization, local topology, and surrounding sequences.

## Materials and Methods

### Growth conditions

*Bacillus subtilis* cells were grown in LB medium for strain constructions and experiments. The antibiotics tetracycline (12.5 μg/ml) and spectinomycin (100 μg/ml) were used as indicated. Growth was monitor by optical density at 600 nm (OD_600_).

### Strains and alleles

The *B. subtilis* strains used were all derived from JMA222 (*trpC2, pheA1*, ICE*Bs1*^0^), a derivative of JH642 (Perego *et al*., 1988; Smith *et al*., 2014) that is cured of ICE*Bs1* (Auchtung *et al*., 2005b). ELC38 (Δ*comK*::*spc*), ELC42 (Δ*rok* Δ*comK*::*spc*), and ELC43 {*att*(*yufKL*)::Tn*916* Δ*comK*::*mls*} were made by natural transformation (Harwood and Cutting, 1990) and are described in more detail below. The *comK* mutants were used to prevent cells from acquiring DNA by transformation rather than conjugation during mating assays. Additionally, loss of *comK* suppresses the growth defect of *rok* mutants (Hoa *et al*., 2002).

The Tn*916* donor strain (ELC43) is derived from CMJ253 (Johnson and Grossman, 2014) and contains a single copy of Tn*916* inserted at the beginning of *yufK*. CMJ253 was generated by natural transformation of JMA222 with genomic DNA from BS49 (Haraldsen and Sonenshein, 2003; Johnson and Grossman, 2014; Browne *et al*., 2015; Wright and Grossman, 2016) and selecting for tetracycline resistance, indicating Tn*916* acquisition. The *comK*::*mls* allele in ELS43 is a deletion-insertion that was generated by replacing most of the *comK* open reading frame (from 47 bp upstream of *comK* to 19 bp upstream of its stop codon) with the resistance cassette from pDG795 (macrolide–lincosamide–streptogramin (MLS)). The cassette was combined with up- and downstream homology arms via isothermal assembly (Gibson *et al*., 2009).

The Δ*comK*::*spc* allele in ELC38 and ELC42 was previously described (Luttinger *et al*., 1996; Auchtung *et al*., 2005a). The *Δrok* allele in ELC42 is an unmarked deletion of *rok* (from 1 bp upstream of *rok* to 11 bp upstream of its stop codon). First, *rok* was replaced with a chloramphenicol resistance cassette (*cat*) flanked by *lox* sites. *cat* was then removed from the chromosome by Cre-mediated recombination between the *lox* sites. *cre* was introduced by transforming cells with pDR244, a temperature-sensitive plasmid that expresses Cre, leaving an unmarked deletion with a small scar (Meisner *et al*., 2013; Johnson and Grossman, 2014). The plasmid was cured by growth at non-permissive temperature.

### Generation of Tn*916* insertion libraries and mating assays

Libraries of Tn*916* insertions were made in both wild type (ELC38) and Δ*rok* (ELC42) *B. subtilis* strains. Briefly, the Tn*916* donor strain (ELC43; has Tn*916* at the beginning of the *yufK* ORF) and recipients were grown in LB medium to mid- or late-exponential phase (OD_600_ of 0.5 and 1, respectively), then mixed at a 1:1 ratio (4 total OD_600_ units of cells) and applied to a nitrocellulose filter. Filters were placed on a 1.5% agar plate with 1x Spizizen’s salts (Harwood and Cutting, 1990) for 3 h at 37°C. Cells were harvested off the filters and plated onto 1.5% agar plates containing tetracycline and spectinomycin to select for transconjugants to be used for insertion sequence analysis. Transconjugants from multiple matings were recovered from petri plates, pooled, and then grown in LB medium to stationary phase before harvesting genomic DNA to map Tn*916* insertion sites by high throughput sequencing. In total we isolated approximately 10^5^ independent transconjugants in each of *rok*+ and *Δrok* strains. Conjugation efficiency (transconjugants per donor) is defined as the number of transconjugant CFUs obtained after mating per donor CFUs added to the mating mixture x 100%.

### Harvesting transconjugant gDNA

Transconjugant colonies were resuspended from tetracycline/spectinomycin agar plates and used to inoculate a culture of LB medium containing 100 μg/ml spectinomycin (to prevent growth of donors). Tetracycline was excluded during growth so as not to stimulate Tn*916* excision from original integration sites (Su *et al*., 1992; Showsh and Andrews, 1992; Manganelli *et al*., 1995; Celli and Trieu-Cuot, 1998; Wright and Grossman, 2016). Theoretically, any residual recipient cells that did not receive a copy of Tn*916* could also grow in the absence of tetracycline, but these should not contribute to our Tn*916*-specific sequencing. Cells were grown to stationary phase (OD_600_ ∼2-3) before harvesting by centrifugation. gDNA was prepared using a Qiagen 100 G tip purification kit. The isolated gDNA had an *oriC/terC* ratio of ∼1, determined as previously described (Anderson *et al*., 2022), indicating that the measured frequencies of insertions in each site was not artificially affected by copy number variation due to ongoing DNA replication.

### DNA sequencing

In general, the <u>Tra</u>nsposon-<u>D</u>irected <u>I</u>nsertion <u>S</u>ite (TraDIS) sequencing pipeline was followed to prepare DNA for sequencing (Langridge *et al*., 2009; Potter and Luo, 2010; Mullany *et al*., 2012; Barquist *et al*., 2016). This strategy uses PCR to enrich for Tn*916*-chromosome junctions from randomly sheared gDNA. Approximately 5 μg of gDNA from each library was sheared into ∼300bp fragments with a Covaris ultrasonicator. Sample sizes were confirmed by gel electrophoresis. Ampure SPRI cleanups were routinely used between each step of DNA preparation. The NEBNext End Repair kit was used for blunt-end repair, followed by the NEBNext dA-tailing kit.

A unique adapter (termed “splinkerette”) was generated to aid in the enrichment of transposon-chromosome junctions via PCR (Potter and Luo, 2010; Barquist *et al*., 2016). One strand of this adapter forms a stable hairpin to which a primer cannot anneal during PCR. Instead, a Tn*916*-specific primer must be used for the first round of amplification. In this way, there should be an enrichment of PCR products containing a Tn*916*-chromosome junction rather than chromosomal DNA alone. The so-called “splinkerette” DNA adapter was prepared by annealing oligonucleotides oELC97 (5’- P*CCACTAGTGT CGACACCAGT CTCTAATTTT TTTTTTCAAA AAAA; P* indicates that the 5’ end of the primer is phosphorylated) and oELC98 (5’- CGAAGAGTAA CCGTTGCTAG GAGAGACCGT GGCTGAATGA GACTGGTGTCGACACTAGTGG*T; *T indicates that the final T residue is attached by a phosphorothioate bond). Oligonucleotides were diluted to 200 μM in 1mM tris-HCl pH 8.5 and mixed in equal volumes before placing in a 98°C heat block for 2 minutes, then cooled to room temperature for annealing to occur. T4 DNA ligase (NEB) was used to anneal the adapter to the dA-tailed DNA fragments.

Adapter-ligated DNA was PCR-amplified (19 cycles) with the Tn*916*-specific P5 primer oELC99 (5’- AATGATACGG CGACCACCGA GATCTACAC<u>T GAGTGGTTTTGACCTTGATA</u> <u>AAGTGTGATA</u> <u>AGTCCAGTTT</u> <u>TTATGCG</u>) where the underlined portion anneals within Tn*916*, and the indexed TraDIS P7 primer oELC100.X (5’- CAAGCAGAAG ACGGCATACG AGATXXXXXX CGAAGAGTAA CCGTTGCTAG GAGAGACCG) where XXXXXX is a barcode sequenced used to identify each library after pooling for sequencing. oELC101.X is unable to bind to its target sequence until after the first round of amplification performed by oELC99.

The resulting DNA was pooled and sequenced on a MiSeq (Illumina) to produce 47 bp reads spanning the Tn*916*-chromsome junction using custom sequencing primer oELC101.2 (5’ - TGAGTGGTTT TGACCTTGAT AAAGTGTGAT AAGTCCAGTT TTTATGCGGA TAACTAGATT TTT). oELC102 (5’ - CCACGGTCTC TCCTAGCAAC GGTTACTCTT CG) was used to sequence the attached barcode. The first 10 cycles were run “dark” because the transposon-specific sequencing primer oELC101.2 anneals 10 bp from the end of Tn*916*, resulting in what would be a mono-template across all samples.

### Analyses of DNA sequence reads

#### Mapping reads to the chromosome

In general, the “Bio::TraDIS pipeline” was followed for sequence analysis (Barquist *et al*., 2016). As described above, data were not collected for the first 10 rounds of sequencing because that portion is within Tn*916* and would result in identical reads from all PCR products. The sequence reads thus begin with the coupling sequence (6 bp variable sequence at the left end of Tn*916* that is derived from the donor) and therefore may not match the host target sequence (Caparon and Scott, 1989; Rudy and Scott, 1994; Scott *et al*., 1994a). We followed the approach of Mullany *et al*. (2012) and removed the first 10 bp of each sequencing read prior to mapping to eliminate potential mismatches with the recipient.

Reads were mapped to the AG174 (JH642) genome sequence (Smith *et al*., 2014). The Tn*916* host strains sequenced contained some engineered differences compared to AG174. Both sequenced strains are ICE*Bs1*-cured and Δ*comK*::*spc*, so these regions lack insertions because they were not present in our experiments. The *rok* allele differed in the two sequenced strains, but no insertions in *rok* were seen in the *rok*^+^ strain (ELC38), so this did not create any false positive decreases in Tn916 insertion frequency in ELS42, the Δ*rok* strain.

The “bacteria_tradis” pipeline (Barquist *et al*., 2016) was used to map reads, allowing up to 2 mismatches. Approximately 87% of reads in each condition mapped to the chromosome (∼1.5 million for each pool), and ∼5% of reads mapped to the circularized form of Tn*916*. For the combined data from both wild type and Δ*rok* strains, insertions were observed at a total of 6977 nucleotide positions.

#### Clustering reads into insertion sites

The 6977 nucleotide positions where reads mapped tended to formed clusters at discrete insertion motifs (Fig 1C). We often observed 1) reads mapping to both the top and bottom strand of the chromosome, which reflects insertions at that locus in both orientations, and 2) insertions at multiple closely spaced bases on the same strand, which is presumably because that the polyA and polyT arms of the insertion motif allow for some slippage in how the integrase binds.

We used custom R scripts to cluster reads that were associated with the same insertion site. We combined reads from the wild-type and Δ*rok* samples for this. At insertion sites that had Tn*916* inserted in both orientations, reads on the bottom strand mapped about 14 nucleotides to the left of reads that mapped to the top strand. This apparent 14 bp 3’ overhang is consistent with a 6 bp 5’ overhang when the mapped positions of the reads are corrected for the 10 bp that was removed from the 5’ end prior to mapping to the chromosome. We did a first-pass assignment of reads into clusters by assigning any read that was more than 14 bp downstream of the previous read, regardless of strand, as a new insertion site, and then ran several diagnostics on this set. We found that for insertion sites that had insertions at more than one position on a strand, these insertions typically spanned 2-5 bps. Using these metrics we identified five chromosomal regions that contained closely spaced but distinct insertion sites that were not correctly processed in our automated approach, and resolved those manually, leading to a final set of 1944 insertions site.

The frequency at which these 1944 insertions sites occur varied over six logs, from 2.9 x 10^-7^ (corresponding to sites with just one read) to 0.22 for the site between the *ykyB* and *ykuC* ORFs. The relative frequency of a Tn*916* insertion in each identified site was estimated by the proportion of sequence reads mapping to that site, but it is not possible to accurately estimate insertion frequencies for sites with very few sequence reads. Although these low abundance insertion sites were not analyzed in detail, they are included in the full data set of insertions (Supplemental Table 1).

#### Annotation of insertion sites

We used the center of the insertion site to determine 1) the gene(s) which the insertion site was within or between, 2) whether the insertion site is within a Rok binding region as reported by (Seid *et al*., 2016), and 3) the DNA sequence surrounding the insertion site. For insertion sites with reads on both strands, we assigned the center as the midpoint between the position with the most reads on top strand and the position with the most reads on the bottom strand. For insertion sites with reads on only one strand, the center was determined by adding or subtracting seven (*i.e.* half the average distance between reads mapping to the top or bottom strand) from the position with the most inserts, depending on whether the reads were mapped to the bottom or top strand, respectively.

### Binding motif and inverted repeats

Binding motifs were determined using MEME version 5.5.6 (Bailey and Elkan, 1994b), which was run online at meme-suite.org. For the general insertion site motif (Fig 3A), we used 61 bp centered on the insertion site from the top 303 insertion sites. The sequences were unweighted, and both strands were searched, allowing for zero or one hit to be found per sequence. A first-order background set from (Zhang *et al*., 2013) was used. To search for inverted repeats, we scanned 61 bp sequences centered on the insertion sites using an online version of Inverted Repeat Finder (Warburton *et al*., 2004). The default settings were used, except we reduced the minimum score to 20.

## Acknowledgements

We thank Stuart Levine and the MIT BioMicro Center for technical assistance and high throughput DNA sequencing. Research reported here is based upon work supported, in part, by the National Institute of General Medical Sciences of the National Institutes of Health under award numbers R01 GM050895, R35 GM122538, and R35 GM148343 to ADG. Any opinions, findings, and conclusions or recommendations expressed in this report are those of the authors and do not necessarily reflect the views of the National Institutes of Health.

## Data availability

Data are available in dataset GSE286917 at NCBI Gene Expression Omnibus database (GEO, http://www.ncbi.nlm.nih.gov/geo).

